# Maternally administered naltrexone and its major active metabolite 6β-naltrexol transport across the placental barrier *in vitro*

**DOI:** 10.1101/2020.04.16.045229

**Authors:** Rajeendra L. Pemathilaka, David E. Reynolds, Nicole N. Hashemi

## Abstract

Opioid use disorder (OUD) has become a growing concern in the U.S. and has been a dominant presence among pregnant women, resulting in an unprecedented amount of prescription medications, particularly naltrexone (NTX), prescribed for pregnant women. Because of unknown potential harm that NTX can impose on the fetus and its premature brain, the needs for safety and regulation of NTX are still undetermined. To address this issue, a microfluidic device is fabricated to mimic structural phenotypes and physiological characteristic of an *in vivo* placental barrier to evaluate near-transport simulations of NTX and its primary metabolite, 6β-naltrexol, across the placental barrier. Following transport analysis, cell layers are evaluated for possible gene-expressions released by an *in vivo* human placenta during NTX and 6β-naltrexol placental exposure. When a 100 ng/mL dose of NTX and 6β-naltrexol (1:1) is administered to the maternal channel, the mean fetal concentration for co-culture models exhibited ~2.5 % of NTX and ~2.2% of 6β-naltrexol of the initial maternal concentration. To prototype and simulate fetal-brain exposure, perfusate from a fetal channel is directed to cultured N27 cells that are then evaluated for gene-expression.

## 1. Introduction

Opioids are composed of natural, artificial, and semisynthetic mediators that can play an essential role in relieving chronic and severe pain,^1^ and the discomfort released by opioid intake has caused a demand upsurge for prescription opioids in the U.S., with more than 650 prescription opioids prescribed every day, particularly for back pain, injuries, and disorders in human body movement.^1,2^ However, this is an alarming increase in opioid intake, with OUD usage more than quadrupling during the past decade.^3^ For example, approximately 115 people fatally overdose from opioids every day, contributing to more than 66% of all drug overdoses,^4,5^ unfortunately including much OUD use by pregnant women.^6^

*In vivo* testing has been the most common form of testing for analyzing NTX concentration levels, with subjects that include mice,^7,8^ rats, ^9^ and rhesus monkeys.^10^ The downside of using these models for testing is that while their anatomies resemble those of human anatomy, their placental tissues lack the analogous cellular and tissue interactions.^11^ In addition to *in vivo* as a form of NTX testing, placebo-controlled testing has also been utilized to some extent.^12–14^ For the most part, these studies have recorded only psychological, and antipruritic reactions from NTX and did not directly evaluate opioid exposure in the placental tissue.^12–14^

The placenta-on-a-chip allows us to co-culture trophoblast and endothelial cells, two of the most prominent cell lines in the human placenta (**Figure 1** (a)-(c)), and to monitor NTX and 6β-naltrexol exposure on the micro-engineered barrier. Previous studies have proven the validity of using placenta-on-a-chip for assessing and replicating different drug transport analysis while mimicking physiological function of human placenta *in vitro*.^15–21^ Through our most recent study, we analyzed caffeine transport across our fabricated microchip, and proved the model to be compatible with withstanding xenobiotic exposure.^21^ Therefore, since placenta-on-a-chip has yet to be tested with opioid contact, our study provides the first appraisal examining NTX and 6β-naltrexol on an organ-on-a-chip.

**Figure 1.**
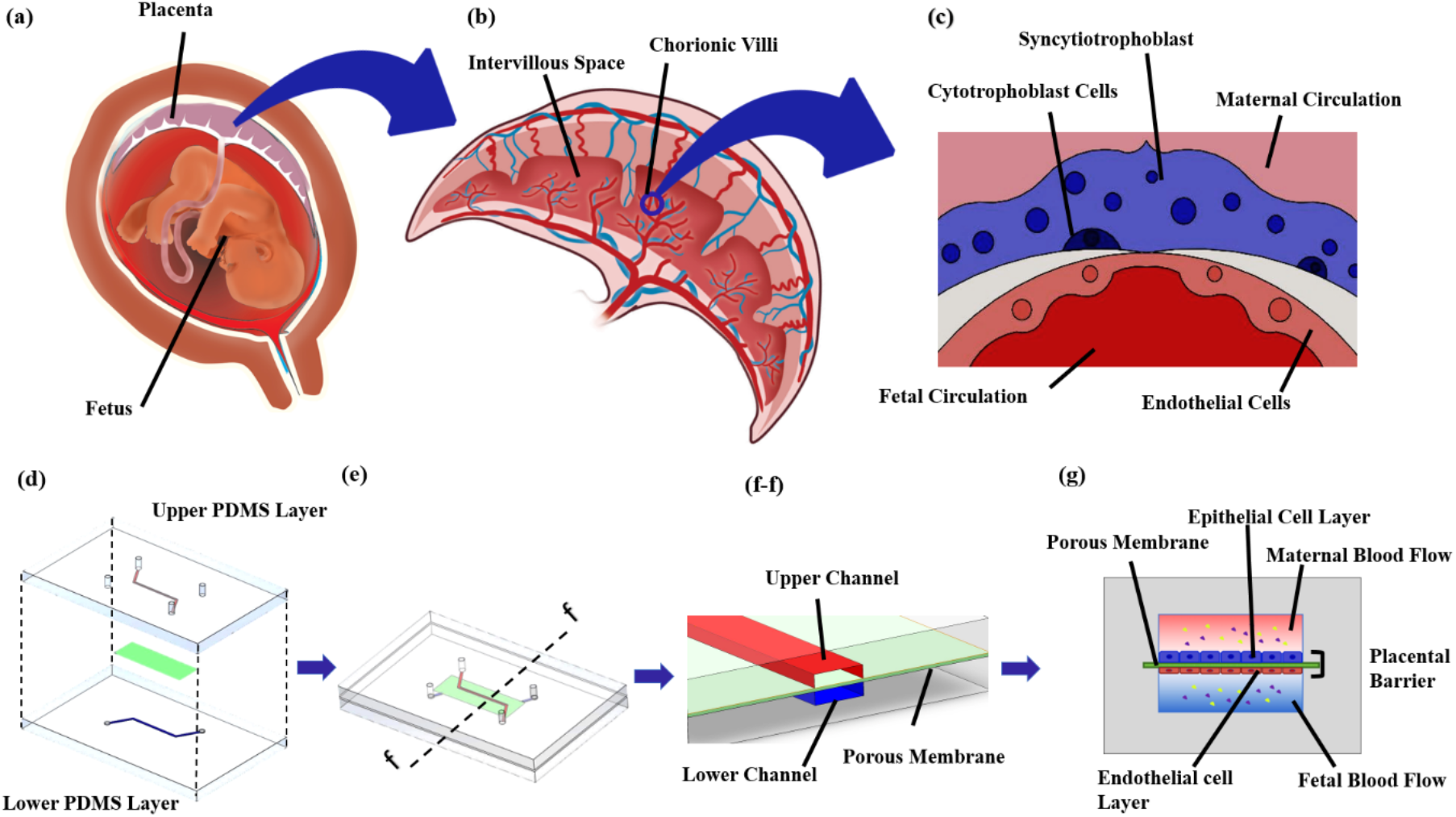
Structure of the human placenta and design of the placenta-on-a-chip device. (a) The placenta is an interim organ that develops only during pregnancy and connects the fetus to the uterine wall via an umbilical cord. (b) Cross-sectional schematic of the human placenta containing important structures called chorionic villi that occupy and demolish uterine decidua and absorb nutritive materials to support fetus maturation; their development is due to the rapid proliferation of trophoblasts. (c) Structure of the placental villi. Syncytiotrophoblast and endothelial cells in maternal and fetal interfaces, respectively, are separated by a basal lamina (a layer of extracellular matrix), at the end of gestational period. During the first trimester, the maternal interface consists of syncytiotrophoblast and cytotrophoblast cells. (d) – (f-f) Exploded, assembled, and cross-sectional views of the fabricated placenta-on-a-chip before microfluidic cell culture. This device, fabricated to resemble the human placental barrier *in vitro,* consists of two microchannel-etched PDMS layers separated by a thin semipermeable membrane. (g) Cross-section of a placenta-on-a-chip after microfluidic cell culture, with the endothelium in the fetal interface and the epithelium in the maternal interface are respectively represented by HUVEC and BeWo cell layers.

This study’s intended purpose was to evaluate opioid transport in our placenta-on-a-chip model, and this stage of testing is made possible through the system and foundation of the placenta-on-a-chip, microfluidic device.^18–21^ Microfluidic devices provide us with the ability to culture and manage cellular and subcellular environments within our microchip, maintain fixed compositions of our cell lines, and replicate fluid movement in parallel layers.^11,22–28^ Our microfluidic device ((Figure 1 (d)-(f-f)) was fabricated using standard lithography techniques.^29–32^ To represent the human placenta’s endothelium and trophoblastic epithelium, human umbilical vein endothelial cells (HUVECs) and trophoblasts cells (BeWo) were cultured accordingly. Once these cells had exhibited behavioral bonding and placental structure *in vitro* after cell seeding and culturing in their respective microchannels, we began exposing our microfluidic device with NTX and 6β-naltrexol. Additionally, by transferring outflow from the fetal channel to cultured N27 embryonic-dopamine cells, we presented a proof-of-concept for modeling NTX and 6β-naltrexol transport to a fetus brain across the placental barrier. We also used a quantitative RT-PCR method to analyze gene-expression changes that ensue from post-exposure to NTX and 6β-naltrexol. This work provides a platform for both scientific and pharmaceutical communities to examine and extrapolate information regarding safety and potential side-effects caused by prescription opioid medication during pregnancy.

## 2. Results and discussion

### 2.1 Fabrication and verification of placental-barrier-like semipermeable membrane

The co-cultured microfluidic device consisted of three layers (Figure 1 (g)) that replicate a human placenta *in vitro*, similar to that from a previous study.^21^ The top layer, or maternal channel, was composed of BeWo cells that represent the trophoblastic epithelium of the human placenta. Beneath the top layer is a thin semipermeable membrane that imitates the human placental barrier, and adhered to the bottom of the porous membrane is the fetal channel, comprised of HUVECs, to recapitulate the endothelium of a human placenta. To validate that the microarchitecture of the microfluidic device’s soluble microenvironments was represented accurately, cell characterization was performed within the chip. After 24-30 hours of microfluidic cell culture, the cells labeled with CellTracker live cells staining (**Figure 2** (a) and (b)) reflected a proliferation over time of cell populations across the porous membrane in the corresponding channels. The dynamic flow environment in both channels precisely resembled the blood circulation within the maternal and fetal interfaces of the human placenta.

**Figure 2.**
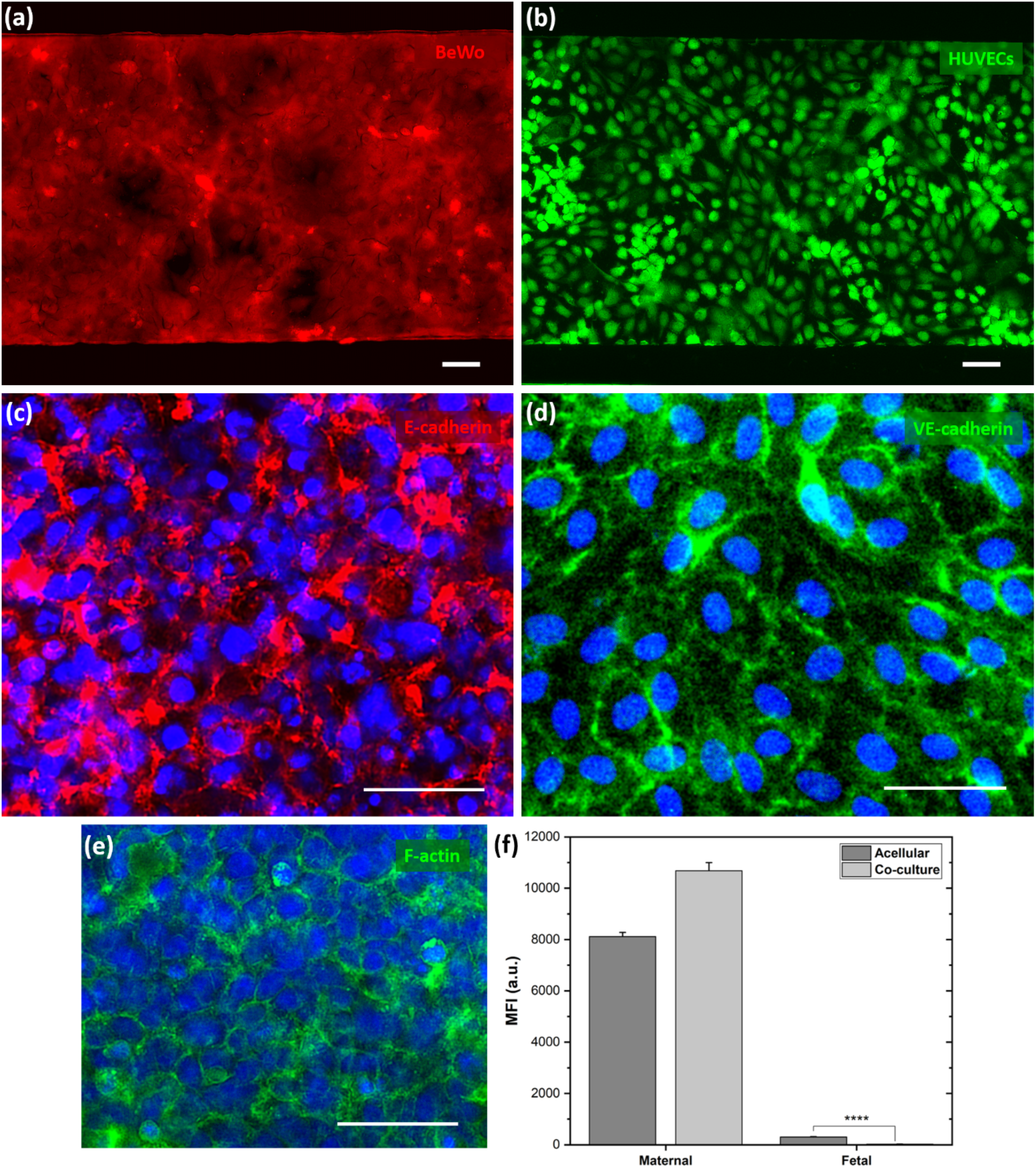
Following 24-30 hours of media perfusion under dynamic flow conditions, epithelial and endothelial cell layers were stained with CellTracker live cell staining. (a) BeWo cells were stained with CellTracker orange and scrutinized for red fluorescent protein (RFP). (b) HUVECs were stained with CellTracker green and analyzed for green fluorescent protein (GFP). (c) Fluorescent microscopic image showing epithelial cells (BeWo) stained with anti-E-cadherin and nuclei labeled with DAPI. E-cadherin is a crucial transmembrane protein to maintain cell-cell junctions in epithelial adherens junctions (AJs). (d) Fluorescent microscopic image displaying endothelial cells (HUVECs) stained with anti-VE-cadherin and nuclei labeled with DAPI. VE-cadherin is an essential endothelial-specific protein responsible for controlling endothelial permeability and maintaining cell-cell junction stabilization in AJs. (e) Formation of microvilli on the apical surface of the trophoblast was assessed by testing for the presence of filamentous actin (F-actin) protein. (f) Placental barrier permeability was investigated using fluorescein-dextran transport across the epithelial-endothelial barrier. *n* = 3 independent experiments. Data represented as mean (± S.E.M.). Scale bars, 50 μm. Two-tailed Student’s t-test, ****, *p* < 0.0001.

After cell proliferation across both microchannels within their corresponding sides of the ECM-coated membrane had been confirmed, after 48 hours of microfluidic cell culture under dynamic flow conditions, we verified the epithelial and endothelial integrity of the microengineered barrier. For investigation of epithelial integrity, E-cadherin antibody was used to detect efficient cell-cell junctions in the epithelium.^33^ VE-cadherin antibody was next used to examine the endothelial integrity with respect to adopting adherens junctions (AJs) in endothelial cells. As shown in Figure 2 (c), strong complexes of E-cadherin detected on the BeWo cells suggested existence of the epithelium cell-cell junctions throughout the membrane in the maternal compartment. VE-cadherin was detected across the HUVECs, validating the existence of a distinct network of AJs on the endothelial cell layer. In addition, the immunofluorescence micrograph in Figure 2 (d) shows that this cell-specific cadherin protein was homogeneously distributed throughout the endothelium, so the complexes displayed from the anti-E-cadherin in epithelium and the anti-VE-cadherin in endothelium confirmed the formation of cell-cell junctions in AJs and the epithelial and endothelial integrity of the placental barrier *in vitro*.

Following the investigation of cell-cell interactions, the maternal channel was evaluated for microvilli formation, because the microvillus plasma membrane is involved in facilitating hormonal and immunological interaction between mother and fetus.^34^ The highly-structured microvilli based in the intervillous space and surrounded by plasma membrane, are considered to be major placental structural components that facilitate materno-fetal transport of nutrients and metabolites through facilitated/simple diffusion, active-transport, phagocytosis, and pinocytosis.^35,36^ As shown in Figure 2 (e), a dense layer of microvilli was observed on the fluorescence microscopy images, and variability this dense layer was detected along the apical surface of the trophoblast cell layer, possibly attributed to differing fluid shear stresses caused by trophoblast cells forming a 3D structure inside the maternal channel, as previously described.^21^ Previous studies for visualizing microvilli under static conditions have resulted in sparse microvillar surfaces.^15,17,19^ In conjunction with such previous studies, microvilli visualized herein dynamic flow conditions have demonstrated elongated microvilli formation in our study.

Thereafter, the placental barrier function was assessed using 3000 MW fluorescein-dextran anionic probes as a permeability assay. Over a 10-hour period, perfusate from both channels was collected from acellular and co-culture devices and analyzed using calculated mean fluorescence intensities (MFIs). As shown in Figure 2 (f), MFIs of the fetal perfusate from acellular devices confirmed a significant passage of fluorescein-dextran (*p* < 0.0001) compared to those collected from the fetal compartment of co-culture devices. Results in Figure 2 (f) also revealed that in co-culture devices, while a few molecules passed through the semipermeable membrane, over a 10-hour period, the overall number of MFIs in the fetal compartment of co-culture devices were negligible compared to those in the perfusates collected from the maternal compartments of both acellular and co-culture devices.

Formation of microvilli in the maternal surface was used as an important physiological characteristic in validating the placental barrier *in vitro*. Considering the ability of replicating structural phenotypes and physiological characteristic of *in vivo* placental-barrier, our placental-barrier-like membrane fabricated on a microfluidic device provides an ideal platform for modeling near-transport simulation of nutrients and metabolites of a human placental barrier.

### 2.2 Analysis of NTX and 6β-naltrexol transport across the placental barrier

In this study, we investigated the possibility of mimicking NTX and its primary metabolite, 6β-naltrexol transport, across our fabricated *in vitro* placental barrier. Previous studies have demonstrated that blood naltrexone concentration of ~8 ng/mL has retrieved to a level of 1.1 ng/ mL within 24 hours after digesting one 50 mg tablet,^37^ and ~2-10 ng/mL was identified as a clinically-relevant naltrexone plasma concentration during sustained naltrexone exposure from naltrexone implants.^38,39^ While oral naltrexone accomplice with weak compliance, e.g., for pregnant women with opioid dependence, long-acting injection of naltrexone provides a potential for using it as a probable medication.^40,41^ Since this study was carried out to demonstrate the possibility of utilizing our placental barrier in a microfluidic device to study the transport of NTX and 6β-naltrexol across human placenta *in vitro*, we introduced a final concentration of 100 ng/mL NTX and 6β-naltrexol (1:1) to the maternal channel. A concentration of 100 ng/mL in this proof-of-concept was also used to receive a detectable level of NTX and 6β-naltrexol concentrations via LC/MS in perfusate collected from the fetal channel.

Initially, as a control condition, we administered NTX and 6β-naltrexol through an acellular device with no epithelial and endothelial cell layers. As shown in **Figure 3** (a), fetal naltrexone concentrations calculated with compiling calibration standards (Figure 3 (b)) exhibited multiple fluctuation phases over a span of 8 hours. NTX concentrations exhibited sudden drops at 4, 6.5, and 8 hours, and between t = 1 to t = 3.5 hours, t = 4 to t = 6 hours, and t = 6.5 to t = 7.5 hours, with similar patterns of increase observed during the study. This could be attributed to active transport of NTX back and forth between maternal-fetal interfaces. Overall, fetal concentration in acellular devices began to rise from an initial value (0.82 ± 0.04 ng/mL) and reached maximum concentrations of 16.88 ± 0.89 ng/mL at 7.5 hours. Conversely, fetal naltrexone concentrations in the perfusate collected from co-culture devices remained nearly stable without major significant fluctuations compared to concentrations in perfusates collected from acellular devices. A slight drop in concentration was observed at 4 hours, and from t = 1 to t = 3.5 hours and t = 4 to t = 8 hours, fetal NTX concentrations maintained a mean of 1.49 ± 0.32 and 3.18 ± 0.31 ng/mL, respectively. Additionally, before and after the slight drop, fetal NTX levels achieved a maximum concentrations of 2.71 ± 1.47 and 4.49 ± 0.83 ng/mL at 3.5 and 7.5 hours, respectively. Overall, the co-culture model maintained a mean NTX concentration that is lower by a factor of ~ 4.21 compared to that of the acellular model.

**Figure 3.**
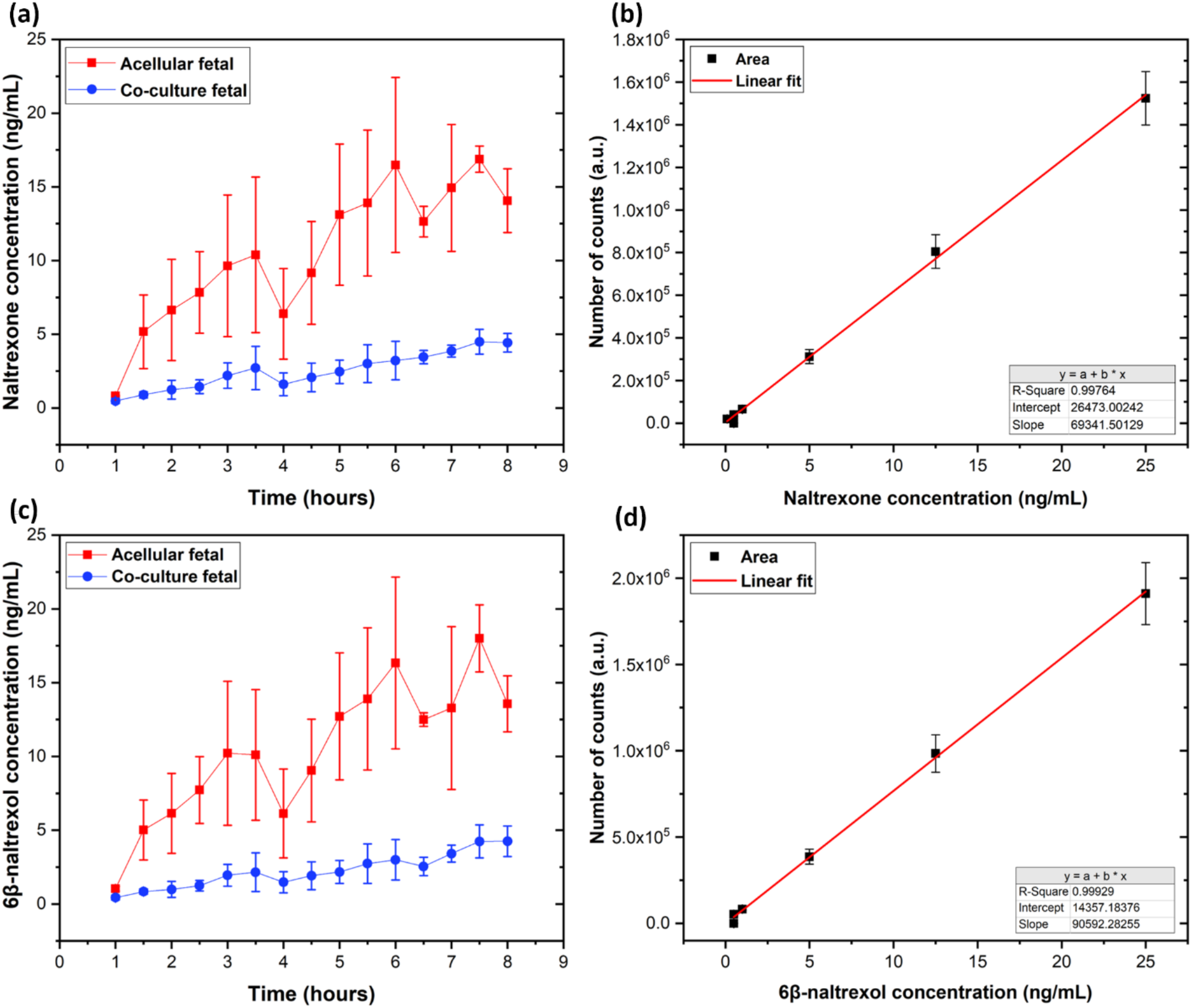
(a) Naltrexone (NTX) concentrations in the fetal channel were quantified via liquid chromatography/mass spectrometry (LC-MS). (b) Calibration standards were plotted to perform NTX analysis. (c) 6β-Naltrexol concentrations in the fetal channel were assessed via LC-MS. (d) Diagram showing calibration standards prepared for quantification of 6β-naltrexol. Co-culture devices have both epithelial and endothelial cells. Acellular devices contain only the bare membrane with perfusion of media as a control condition. n = 3 independent experiments. Data represented as mean (± S.E.M.).

Similarly, fetal 6β-naltrexol concentrations were analyzed in perfusates collected in acellular and co-culture devices. As shown in Figure 3 (c), final 6β-naltrexol concentrations were quantified after interpolating with generated concentration standards (Figure 3 (d)). Correspondingly to fetal NTX concentrations in acellular devices, fetal 6β-naltrexol concentrations in acellular devices exhibited similar trend over a span of 8 hours. The concentrations of 6β-naltrexol in acellular devices exhibited rapid decrease at 4 and 6.5 hours, and, in contrast to fetal NTX concentration in coculture devices, this trend was also observed in fetal 6β-naltrexol concentrations in co-culture devices. For those hours, concentrations of 6.13 ± 3.01 and 12.50 ± 0.46 ng/mL for acellular devices and 1.47 ± 0.72 and 2.54 ± 0.62 ng/mL for co-culture devices were detected, respectively. In comparison, mean fetal 6β-naltrexol concentrations from t = 1 to t = 3.5 hours, t = 4 to t = 6 hours, and t = 6.5 to t = 8 hours were observed as 6.71 ± 1.329, 13.00 ± 2.15, and 14.34 ± 1.49 ng/mL for acellular devices and 1.27 ± 0.28, 2.26 ± 0.43, and 3.61 ± 0.43 ng/mL for co-culture devices, respectively. The mean fetal 6β-naltrexol concentrations that were analyzed from perfusates collected from acellular devices exhibited a 4.67-fold higher level those in co-culture devices, for an interval of 8 hours.

Interestingly, in comparison with the initially-administered maternal NTX and 6β-naltrexol concentration (100 ng/mL), the mean fetal NTX and 6β-naltrexol concentrations evinced ~2.5% and ~2.2% levels in devices in the presence of HUVECs and BeWo cells, respectively. Conversely, in the absence of HUVECs and BeWo cells, NTX and 6β-naltrexol concentrations were ~10.5% and ~10.3% levels with respect to the initial maternal concentrations. Uncommonly, the mean NTX and 6β-naltrexol concentrations evaluated for co-culture model began to rise after 6 hours and continued to increase until the end of the experiments, reaching higher mean concentrations compared to those from t = 1 to t = 6 hours (NTX: 1.94 ± 0.27; 6β-naltrexol: 1.72 ± 0.26). After 6 hours, concentrations sustained mean concentrations of 4.06 ± 0.29 and 3.61 ± 0.43 ng/mL for NTX and 6β-naltrexol, respectively. We suspect that this may be due to a disturbance of barrier integrity of the epithelial and endothelial cell layers in our microfluidic device. If there was such a possible disruption in HUVECs and BeWo cells layers, a rise in concentration levels would be expected for both NTX and 6β-naltrexol because structural and physiological functions were not precisely represented during this scenario. Initially, to verify whether there was a disruption in epithelial and endothelial cell layers, after 8 hours, BeWo cells and HUVECs in microchannels were stained with CellTracker to visualize live cells, and as a control condition, the maternal channel in co-culture devices was perfused with F-12K medium with the absence of NTX and 6β-naltrexol. As shown in **Figure 4** (a), for control conditions, the epithelial cell layer was not disrupted as seen for the co-culture devices with exposure to NTX and 6β-naltrexol ((Figure 4 (b)), and similar observations were identified for the endothelial layer. HUVECs stained for live cells in the control ((Figure 4 (c)) appear to have a matured cell layer, compared to the raptured cell layer in the co-culture devices with perfusing NTX and 6β-naltrexol ((Figure 4 (d)).

**Figure 4.**
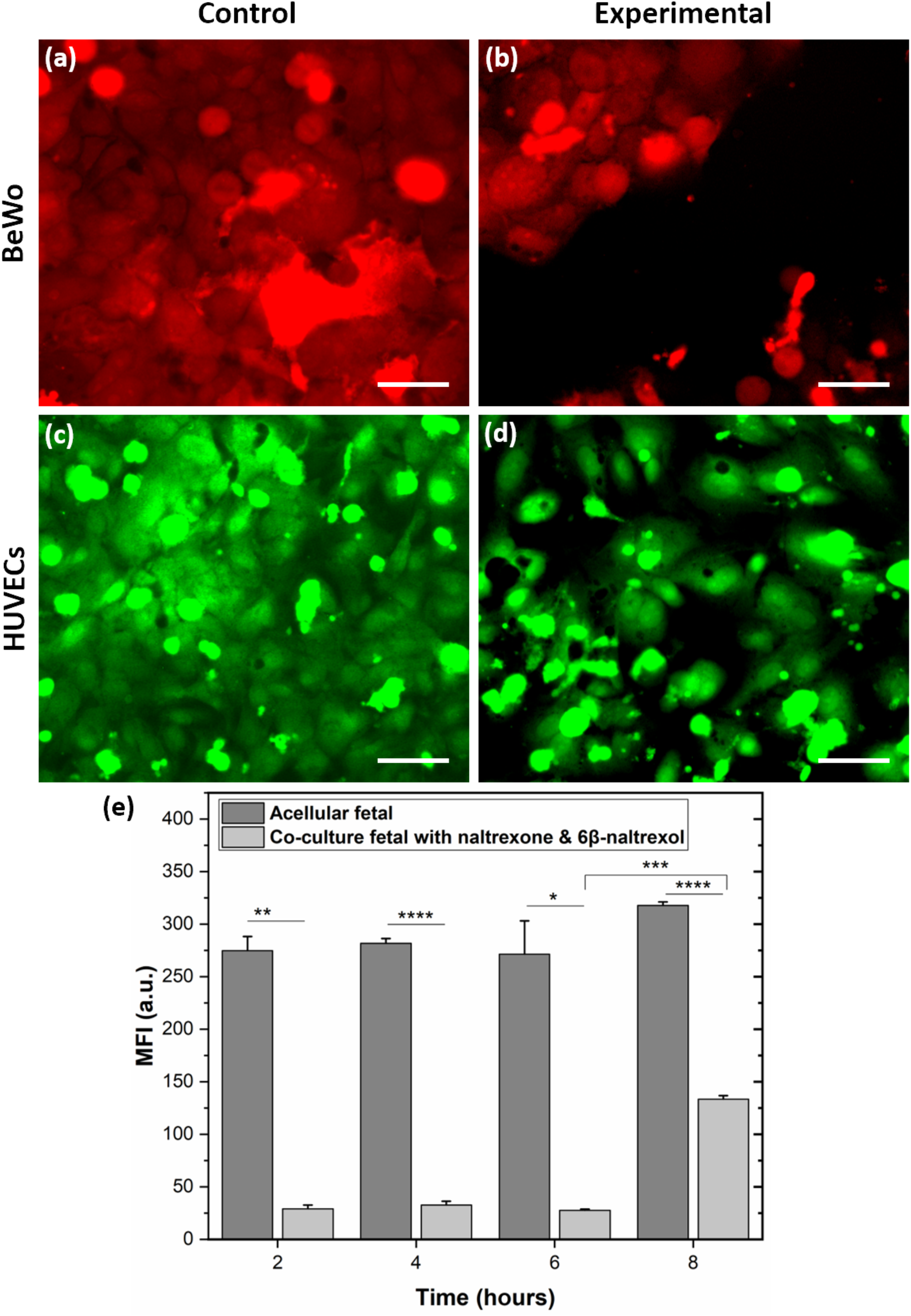
Following NTX and 6β-naltrexol exposure, epithelial and endothelial cell layers were stained with CellTracker live-cell staining. Control and experimental conditions represent co-culture devices perfused with and without NTX/6β-naltrexol in the maternal flow, respectively. BeWo cells and HUVECs were stained with CellTracker orange and green, respectively. (a) BeWo cell layer under control conditions. (b) BeWo cell layer under experimental conditions. (c) HUVEC cell layer under control conditions. (d) HUVEC cell layer under experimental conditions. (e) Transport analysis of fluorescein-dextran across acellular and co-culture devices to evaluate barrier permeability during the NTX/6β-naltrexol transport study. n = 3 independent experiments. Data represented as mean (± S.E.M.). Scale bars, 50 μm. Two-tailed Student’s t-test, *, p < 0.05; **, p < 0.01; ***, p < 0.001; ****, p < 0.0001.

Barrier permeability function was also evaluated to identify the time frame in which the disruption of cell layers occurs on the fabricated placental membrane. As shown in Figure 4 (e), no significant differences were observed, and virtually-constant levels of MFIs were displayed in perfusates collected from co-culture devices during the first six hours. The MFIs evaluated from t = 6 to t = 8 hours in co-culture devices implied a statistical-significant increase (*p* < 0.001) compared to MFI at t = 6 hours. In comparison with MFI levels measured over 6 hours in co-culture devices, the levels showed ~4.5 times greater values in perfusates evaluated from t = 6 to t = 8 hours. Interestingly, no significant differences were observed within MFIs of perfusates collected from acellular devices at 2-hour intervals for 8 hours. In addition, the MFIs in co-culture devices always remained lower than those evaluated in perfusates collected from acellular devices. This not only verified that the disruption visualized in microscopic imaging of HUVEC and BeWo cell layers occurred between t = 6 to t = 8 hours, but also confirmed that, after 6 hours of NTX/6β-naltrexol exposure, the *in vitro* placental barrier became profoundly permeable to fluorescein-dextran. This placental barrier alteration existing after 6 hours of NTX and 6β-naltrexol perfusion also supported the possibility of becoming eminently permeable to aforesaid drugs. Such a rupture of the *in vitro* placental barrier could be attributed to cell apoptosis due to effects of NTX and 6β-naltrexol on epithelial and endothelial cell layers.

### 2.3 Gene-expression analysis of the placental barrier following post-NTX/6β-naltrexol exposure

In this phase of the study, we evaluated possible genetic changes that the placental barrier undergoes following post-NTX and −6β-naltrexol exposure. A previous study reported that low-dose naltrexone (LDN) exhibited a reduced plasma concentration of interleukin (IL)- 6 and tumor necrosis factor (TNF)-α.^42^ TNF-α is considered a proinflammatory cytokine that worsens disease,^43^ and IL-1α is another proinflammatory cytokine that binds IL-1 receptor.^44^ Conversely, IL-6 not only performed as a proinflammatory cytokine, but also an anti-inflammatory myokine and a cytokine involved in responding to inflammation and infection.^45,46^ IL-8 is another cytokine responsible for activating neutrophils in inflammatory regions.^47^ These properties were evaluated with IL-1α, IL-6, and TNF-α for the HUVECs layer and IL-1α, IL-6, and IL-8 for the BeWo cell layer. As shown in **Figure 5** (a), proinflammatory cytokines, IL-1α and IL-6, implied lower fold changes in gene-expression levels when exposed to NTX and 6β-naltrexol compared to levels reported under control conditions. Interestingly, only IL-1α exhibited a significant reduction (*p* < 0.01) even though IL-6 also exhibited a decrease in fold change. TNF-α gene-expression also revealed an increase in fold change levels, but no significant difference was detected. Intriguingly, when exposed to NTX and 6β-naltrexol, IL-1α and IL-6 fold change levels in HUVECs (Figure 5 (b)) exhibited completely different tendencies compared to fold change levels achieved in BeWo cells. When evaluated for IL-1α and IL-6 in HUVECs, fold change displayed higher levels with no significant increase than those measured under control conditions. This change in fold change patterns could be attributed to epithelial and endothelial cell apoptosis during NTX/6β-naltrexol exposure. IL-8, a cytokine responsible for activating neutrophils in inflammatory regions, and evaluated in the endothelial cell layer, exhibited a decrease in fold change compare to that under control conditions. Since no significant difference was observed, IL-8 presented the possibility of achieving inflammatory-free conditions in fetal interfaces during an NTX/6β-naltrexol exposure. Due to the lack of availability gene-expression analysis performed on the maternal and fetal interfaces during an NTX/6β-naltrexol exposure, further investigation is recommended to validate these results before comparing them to gene-expression analysis of *in vivo* human placental barrier.

**Figure 5.**
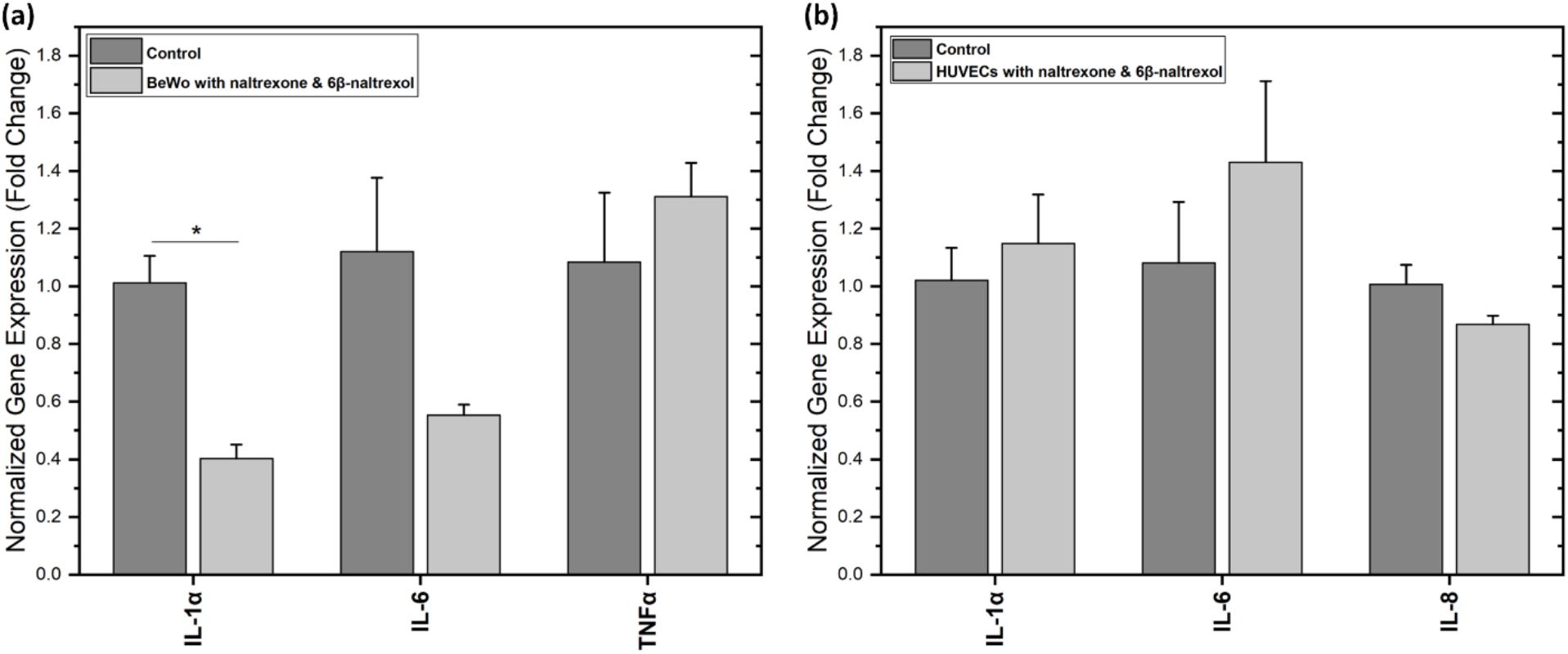
Gene-expression analysis of epithelial and endothelial cells following NTX and 6β-naltrexol exposure, via RT-PCR method. Control conditions consisted of cells in a co-culture device with perfusing NTX- and 6β-naltrexol-free media through the maternal channel. Expression levels were reported as a fold change in gene-expressions analogous to that of human 18S rRNA, the housekeeping gene. (a) BeWo cells from maternal side were assessed for IL-1α, IL-6, and IL-8 genes. (b) HUVECs from the fetal side were evaluated for IL-1α, IL-6, and TNF-α genes. n = 3 independent experiments. Data represented as mean (± S.E.M.). Two-tailed Student’s t-test, *, p < 0.01.

### 2.4 N27 embryonic-dopamine cell line exposed to NTX and 6β-naltrexol

In the next phase of the study, we proposed a possible concept for simulating effects on fetus brain cells from maternally-administered NTX and 6β-naltrexol to the placental barrier, because a previous study on pregnant mice reported that 6β-naltrexol enters the fetal brain at greater levels after promptly crossing the placental barrier.^48^ Initially, N27 cells, plated and cultured for 5 days, were exposed to NTX and 6β-naltrexol by simply directing the outflow of the fetal channel, and two control condition measurements were also used to compare the results obtained from N27 cells exposed to NTX and 6β-naltrexol. Following an 8-hour transfer of perfusate to N27 cells cultured in 6-well plates, cells were stained with a live/dead cell assay. As indicated in **Figure 6** (a) and (b), fluorescent images displayed only minor cell death under both control conditions, while live/dead cell assays identified a large amount of cell death on the cell layer (Figure 6 (c)) exposed to the fetal channel from co-culture devices perfusing NTX and 6β-naltrexol. To quantify cell viability, fluorescent images were analyzed by counting live and dead cells. Cell viability (Figure 6 (d)) exhibited no significant difference (*p* > 0.05) for both control studies, verifying minimal effects against cell apoptosis after N27 cells exposed to EGM. In addition, an experimental study with N27 cells exposed to NTX and 6β-naltrexol showed a significant decrease in cell viability compared to cell viability found in both control studies (*p* < 0.001).

**Figure 6.**
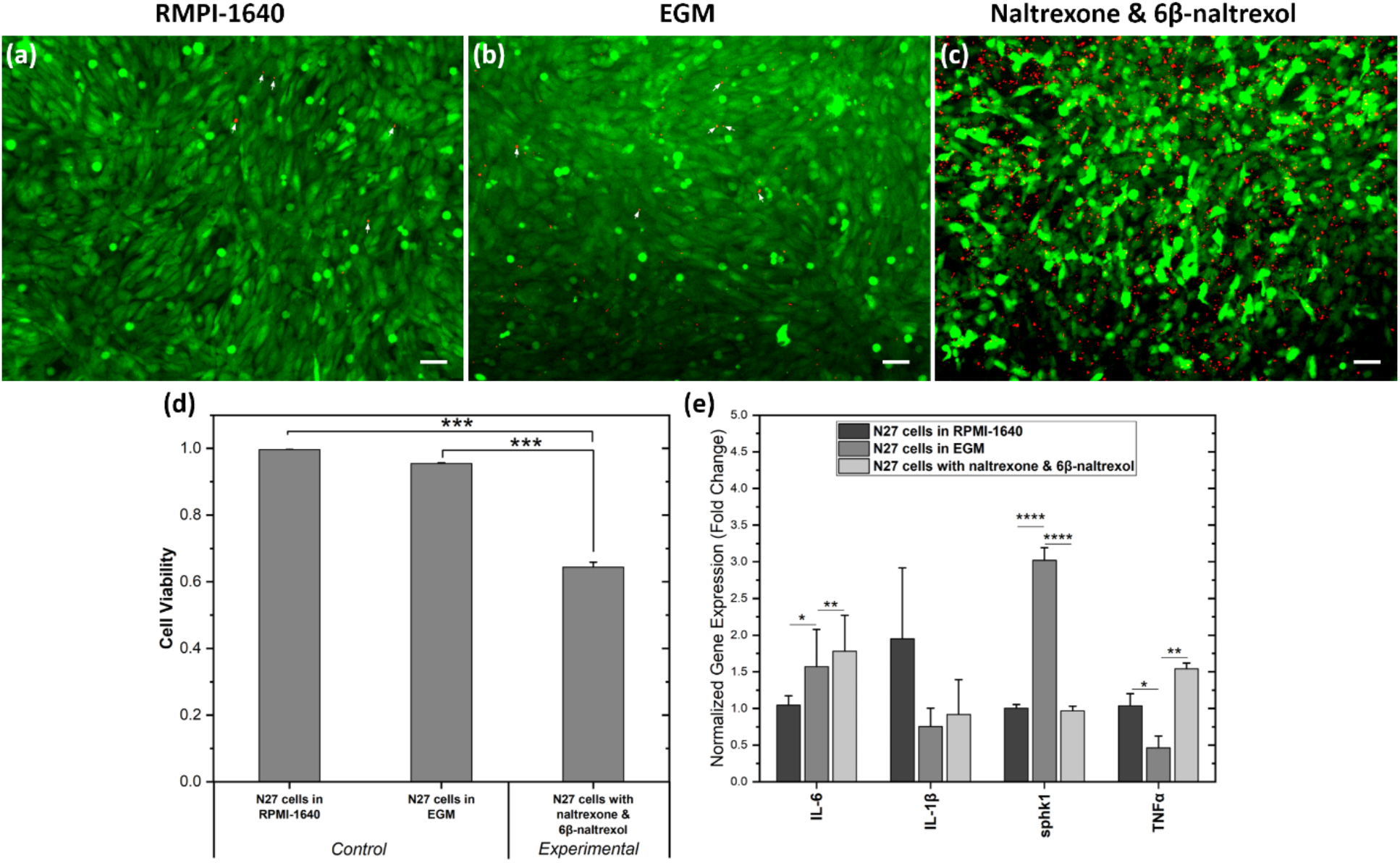
The outflow from the fetal channel was directed toward N27 cells cultured in 6-well plates and a live/dead assay was performed on the cells after 8 hours of continuous exposure to the following conditions: (a) Cells were maintained in RPMI 1640 medium with no exposure to outflow from the fetal channel. (b) Cells (in RPMI 1640 medium) were exposed to the fetal-channel outflow from the co-culture devices that perfused NTX- and 6β-naltrexol-free medium through the maternal channel (c) Cells (in RPMI 1640 medium) were exposed to NTX and 6β- naltrexol through the fetal-channel outflow from co-culture devices. Live and dead cells indicated in green and red, respectively. (d) N27 cells under same conditions as above quantified to examine cell viability. Data were calculated from 3-4 images. (e) Gene-expression analysis on N27 cells subjected to conditions (a), (b), and (c). The mouse 18S rRNA, the housekeeping gene, was referenced to report gene-expression levels as a fold change. n = 3 independent experiments. Data represented as mean (± S.E.M.). Scale bars, 50 μm. One-way ANOVA, *, p < 0.05; **, p < 0.01; ***, p < 0.001; ****, p < 0.0001.

We next evaluated N27 cells for genetic changes following post-exposure to NTX and 6β-naltrexol. It has been reported that IL-6 and IL-1β expression levels have produced increases in plasma levels of fetal brains,^49^ and acute inflammatory insult to a developing brain from IL-6 gene-expression,^50^ and the possibility of TNF-α reducing embryonic development of the brain have also been reported.^51^ Sphingosine kinase (sphk)1 enzyme is associated with increasing survival and proliferation of cells,^52^ and sphk1 exhibits standard physiological functions in developing brain cells.^53^ As indicated in Figure 6 (e), N27 cells exposed to NTX and 6β-naltrexol exhibited significantly higher fold change levels in IL-6 compared to those under both control conditions, while the control conditions exhibited a significant difference (*p* < 0.05) in fold change levels for IL-6 gene-expression. Fold change levels revealed lower expression levels of IL-1β in N27 cells exposed to EGM and NTX/6β-naltrexol compared to levels in N27 cells maintained in RPMI-1640, but no significant differences were observed. Interestingly, sphk1 gene-expressed fold change levels measured in N27 cells in RPMI-1640 and N27 cells exposed to NTX and 6β-naltrexol remained virtually-constant while the levels for cells exposed to EGM showed a significantly higher fold change values compared to cells in RMPI-1640 and exposed to NTX and 6β-naltrexol (for both, *p* < 0.0001). This could be attributed to rapid cell growth in N27 cells when exposed to EGM, because extra growth factors in EGM could be promoting cell growth. Conversely, TNF-α gene-expression exhibited significantly lower fold change values in N27 cells in RMPI-1640 were observed than in N27 cells exposed to NTX and its primary metabolite, and EGM. Further studies are warranted to validate these results achieved from gene-expression analysis of N27 cells following post-exposure to NTX and 6β-naltrexol.

## 3. Conclusion

In our study, we fabricated a human placental barrier *in vitro*, allowing us to investigate NTX and 6β-naltrexol transport across our micro-engineered barrier. Initially, *in vitro* placental barrier was evaluated for structural phenotypes and physiological characteristics of a human placenta. Following the barrier verification, 100 ng/mL of NTX and 6β-naltrexol was introduced to the maternal channel and perfused for 8 hours, and mean fetal NTX and 6β-naltrexol concentrations over this interval were recorded as 2.50 ± 0.26 and 2.22 ± 0.25 ng/mL, respectively, for co-culture devices. The epithelial cell layer after NTX and 6β-naltrexol exposure was evaluated for IL-1α, IL-6, and TNFα and endothelial cell layer was examined for IL-1α, IL-6, and IL-8 genes. During the next phase of the study, perfusate for the fetal channel was directed to investigate embryonic brain cells exposed to NTX and 6β-naltrexol. Following cell viability evaluation, cells were observed for IL-6, IL-1β, sphk1, and TNFα gene-expressions. With enhanced detection through LC-MS, this proof-of-concept can be used to analyze the transport of ~2-10 ng/mL (clinically-relevant plasma concentration for NTX) of NTX and 6β-naltrexol and its effects on a fetus and its premature brain.

## 4. Methods

### 4.1 Cell culture

HUVECs and BeWo were chosen to present the cells at the fetal and maternal interfaces, respectively, as previously described.^21^ BeWo cells (ATCC) were cultured in Kaighn’s Modification of Ham’s F-12K medium (Thermofisher), supplemented with growth factors and 10% fetal bovine serum (FBS, Gibco). HUVECs (Lonza) were cultured in endothelial growth medium (EGM, R&D Systems). N27 embryonic-dopamine cell line was generously provided by Dr. Anumantha Kanthasamy at Iowa State University. N27 cells were cultured in RPMI 1640 medium (Gibco), supplemented with 10% FBS, 2mM L-glutamine (Gibco), and 100 U/mL penicillin and streptomycin (Gibco). Cell lines were incubated at 37 °C with 5% CO_2_ in air until they were 80-90% confluent. After at least three passages, N27 cells were dissociated with trypsin/EDTA (1X) (Cascade Biologics) and plated into individual wells of a 6-well plate at a density of 25 ×10^3^ cells/well.

### 4.2 Design and fabrication of the chip

The placenta-on-a-chip consists of two polydimethylsiloxane (PDMS) slabs, each with a microchannel constrained by the following dimensions: 100 μm (height) and 400 μm (width). The microfluidic device was fabricated using a silicon wafer SU-8 mold produced through standard soft lithography techniques. A mixture of PDMS base and curing agent solution (Dow Corning) at 10:1 (w/w) was introduced into a SU-8 mold placed in a petri dish (15 cm in diameter), as previously described.^21^ The PDMS was cut and removed from the mold as upper and lower layers after the PDMS had solidified at room temperature. A biopsy punch was then used to produce inlet/outlet holes (1 mm in diameter) on the upper PDMS layer. A polyethylene terephthalate (PET) membrane (0.4 μm pore size) as the barrier between two channels, taken from the membrane inserts (Corning®), was placed over the mid-section of the lower channel, after which both PDMS layers were plasma-treated, aligned, bonded together, and left overnight to completely cure the bond. The chip was UV-sterilized for 20 minutes following attachment of inlet/outlet tubing to provide fluid access to each channel, as described previously.^21^

### 4.3 Microfluidic cell culture

Following UV-sterilization, an Entactin-collagen IV-laminin (E-C-L, Millipore) solution (10 μg/mL in sterile serum-free medium) was coated across both faces of the PETE membrane through the channel inlets, and the chip then was refrigerated overnight at 4°C. On the following day, phosphate-buffed saline (PBS) was perfused twice to remove and wash excess E-C-L from the microfluidic device’s channels, after which the microchip was prepared for microfluidic cell culture. Before infusion, both cell lines were dissociated with trypsin/EDTA (1X) and cells were resuspended in full growth medium (EGM or F-12K medium). Initially, the HUVECs were introduced into the lower channel at a density of ~5 × 10^6^ cells/mL, then incubated at an inverted position at 37°C with 5% CO_2_ in air for 1 hour to ensure adherence of cells to the membrane. Similarly, the BeWo cells were seeded into the upper channel at a density of ~5 × 10^6^ cells/mL, and after the cells had exhibited adhesion to the membrane, 3 mL syringes (Becton, Dickinson, and Company) filled with EGM and F-12K medium were connected to the inlets of the lower and upper channels, respectively. The syringes were then connected to a syringe pump operated at a flow rate of 50 μL/hour.

### 4.4 Live cell staining

The BeWo cells and HUVECs were initially stained with CellTracker orange and green (Life Technologies), respectively. Both cell lines were incubated at 37°C with 5% CO_2_ for 45 minutes with staining in a diluted serum-free medium before dissociation, conforming to the manufacturer’s recommended protocol.

### 4.5 Immunohistochemistry

Investigation of the epithelial and endothelial integrity was carried out after confirming proliferation of cells in the microchannels. Initially, cultures in the fetal and maternal channels were fixed in 4% paraformaldehyde at room temperature for 20 minutes then rinsed with PBS, after which the microchannels were incubated with a blocking buffer (0.4% bovine serum albumin (BSA), 0.2% Triton X-100, and 5% normal donkey serum (Jackson Immuno Research Labs)) for 60 minutes at room temperature. After incubation, microchannels were incubated overnight at 4°C with anti-vascular endothelial-cadherin (VE-cadherin) and anti-epithelial cadherin (E-cadherin) (Cell Signaling Technologies) and primary antibodies were diluted with blocking buffer for HUVECs and BeWo cells, respectively. On the following day, the microchannels were rinsed with PBS followed by incubation with secondary antibodies and DAPI solution (diluted with blocking serum) in the dark for 90 minutes. Finally, both microchannels were rinsed with PBS and the chip’s membrane was carefully separated from the microchip, mounted to a coverslip, and examined for immunostaining using an inverted fluorescence microscope (Zeiss Axio Observer Z1).

### 4.6 Staining to visualize microvilli formation

After 72 hours of proliferation of cells in the channels, BeWo cells were stained for filamentous actin (F-actin) to visualize formation of microvilli. Following a PBS rinse, the maternal channel was fixed with 4% paraformaldehyde for 15 minutes, after which the cultures in the channel were rinsed with PBS and permeabilized with 0.5 % Triton X-100 for 10 minutes at room temperature. After another PBS rinse, cultures were stained with CF® 488A-conjugated phalloidin (Biotium) (5 μL of a 200 U/mL (in cell culture grade water) stock solution in 200 μL of PBS). Following a counterstain of cell nuclei with DAPI, cells were incubated for 20 minutes at room temperature, washed with PBS, then imaged with a Zeiss Axio Observer Z1 Microscope.

### 4.7 Investigating the barrier permeability

The barrier permeability function of the transport between the fetal and maternal microchannels was analyzed using 3000 MW fluorescein-dextran anionic probes (Invitrogen, ThermoFisher), as described previously.^21^ The Fluorescein-dextran was initially diluted in PBS to 100 mg/mL, in conformance with the manufacturers’ recommended protocol, then the fluorescein-dextran solution was placed in F-12K medium and brought to a final concentration of 0.1 mg/mL. The maternal channel’s F-12K supplement was then replaced and perfused with dextran-mixed F-12K for 10 hours. At subsequent 2-hour intervals, the perfusate from both the fetal and maternal channels was collected and its fluorescence intensity quantified utilizing a microplate reader (BioTek Synergy 2).

### 4.8 NTX and 6β-naltrexol transport across the placental barrier

NTX and 6β-naltrexol were generously provided by Dr. Wolfgang Sadee and Dr. John Oberdick of Ohio State University. NTX and 6β-naltrexol at 1:1 (w/w) were first diluted to 1 mg/mL (in cell culture grade water) then diluted to a final concentration of 100 ng/mL (in F-12K medium). The medium supplement for the maternal channel was then substituted for the NTX/6β-naltrexol-mixed F-12K medium and perfused for 1 hour to remove the NTX/6β-naltrexol-free F-12K medium from the maternal channel. After 1 hour of equilibrium, perfusate was collected every 30 minutes from the fetal channel and analyzed by liquid chromatography/mass spectrometry (LC-MS), and the permeability between the fetal and maternal channels in the microfluidic device during NTX/6β-naltrexol perfusion was analyzed using dextran anionic probes. Fluorescein-dextran was diluted in NTX/6β-naltrexol-mixed F-12K medium (to a final concentration of 0.1 mg/mL) and perfused through the maternal channel, after which the outflow was sequentially collected at 2-hour intervals and its fluorescence intensity was evaluated using a microplate reader, as described above.

### 4.9 LC-MS for detection of NTX and 6β-naltrexol

An Agilent Technologies 6540 Ultra-High-Definition (UHD) Accurate-Mass Quadrupole Time-of-Flight (Q-TOF) LC-MS system equipped with an Agilent Technologies Eclipse C18 1.8 μm 2.1 mm × 100 mm column (W. M. Keck Metabolomics Laboratory, Iowa State University) was used to perform analysis of NTX and 6β-naltrexol in collected outflow samples. The Agilent system was equipped with a thermostatically-controlled dual-needle injector coupled to an Agilent Technologies 1290 Infinity Binary Ultra High-Pressure Liquid Chromatography (UHPLC) system. The samples were initially transferred into 300 μL conical volume inserts (Thermo Scientific) in 2 mL Surestop vials (Thermo Scientific), loaded onto the Autosampler tray and kept at 4°C during the Q-TOF measurement. A gradient of 1.2 mM ammonia formate (pH 3.5) in High Performance Liquid Chromatography (HPLC) grade water (buffer A) and 1.2 mM ammonia formate (pH 3.5) in a mixture of HPLC grade acetonitrile and methanol (1:2) (buffer B) was used for chromatographic separation. After a 0.2-minute holding time in 100% buffer A, the gradient had transitioned to 33.3% and 48% buffer B in 5.2 to 6.2 minutes, respectively. The composition was switched to 100% buffer B at 8 minutes, followed by a 1-minute hold before returning it to initial conditions by a 4-minute post-run. Electrospray ionization in positive mode was used to detect NTX and 6β-naltrexol as [M + H]^+^ ions (flow rate: 0.4 mL/minute, injection volume: and 10 μL). NTX and 6β-naltrexol were identified at m/z 342.17 and 344.18, respectively. Calibration standards were prepared as a diluted series of combined NTX and 6β-naltrexol at 1:1 (w/w) (first diluted in cell culture grade water then to a final concentration in F-12K medium) at the following concentrations: 0.1. 0.5, 1, 5, 12.5, and 25 ng/mL. Agilent Mass Hunter Quantitative Analysis B.07.00 software was utilized to quantify the detection of NTX and 6β-naltrexol, as in the example of extracted ion chromatograms shown in **Figure S1** and **S2**. The calculated concentrations of NTX and 6β-naltrexol were adjusted to their final values based on relative volume of samples collected from the outflow of the fetal channel.^54,55^

### 4.10 Midbrain-derived embryonic N27 cell exposure to NTX and 6β-naltrexol

After 5 days of culture, the N27 cells in the 6-well plates were prepared for exposure to NTX and 6β-naltrexol. Following the NTX/6β-naltrexol-mixed F-12K perfusion, outflow of the fetal channel was directed to N27 cell-cultured individual wells of a 6-well plate. The treatment was performed for 8 hours after 1 hour of equilibrium phase, as described above. Three different experiment groups including two control conditions were used for data analysis under the following conditions: N27 cells cultured in RPMI 1640 medium and had no exposure to NTX and 6β-naltrexol, N27 cells (in RPMI 1640 medium) exposed to the outflow from the fetal channel without NTX and 6β-naltrexol in system, and N27 cells (in RPMI 1640 medium) treated with NTX and 6β-naltrexol through the outflow of the microfluidic channel, for 8 hours.

### 4.11 Quantitative RT-PCR

After exposure to NTX and 6β-naltrexol, HUVECs and BeWo cells from the microfluidic device or N27 cells from 6-well plates were quantified using the RT-PCR method. After treatment, control and experimental samples were trypsinized, pelleted, and frozen at −80°C, then integrated into single-control and single-experimental sets before homogenization in TRIzol reagent (Invitrogen, ThermoFisher). Following homogenization, RNA isolation and reverse transcription were performed using the Absolutely RNA Miniprep kit (Stratagene) and cDNA synthesis system (Applied Biosystems), respectively. A Qiagen RT^2^ SYBR Green master mix with validated qPCR human primers (for HUVECs and BeWo) or mouse primers (for N27 cells) from Qiagen (Frederick) were used to determine relative magnitudes of gene-expression levels using RT-PCR. Human 18S rRNA or mouse 18S rRNA, the housekeeping genes, were used to normalize each sample, and melting curves and dissociation curves were constructed to verify the gathering of nonspecific amplicons-free peaks, as described in manufacturer’s recommended guidelines. The ΔΔC_t_ method developed to utilize threshold cycle (C_t_) values from housekeeping gene and respective gene was used to calculate and report the results as a fold change in gene-expression.^56,57^

### 4.12 Live/dead cell assay

After exposure to NTX and 6β-naltrexol, assessment of both live and dead cells on the N27 cell line was performed with a combination of 10 μM CellTracker green and 4 μM Propidium Iodide (PI, Invitrogen, ThermoFisher) solutions. Each solution was diluted in serum-free medium, following the manufacturers’ recommended protocol. Live and dead cells were detected with green fluorescent protein (GFP) and red fluorescent protein (RFP), respectively, using an Inverted Microscope (Zeiss Axio Observer Z1). The numbers of live and dead cells were determined using ImageJ software.

### 4.13 Statistical analysis

All experiments were repeated at least three times, with results reported as mean ± S.E.M. Data analysis was performed using a two-tailed Student’s t-test or one-way analysis of variance (ANOVA) using MATLAB (The MathWorks Inc.) or OriginPro (OriginLab Corp.), respectively.

## Author Contributions Statement

R.L.P designed and performed the experiments. R.L.P and D.E.R. analyzed the data and wrote the main manuscript text. The work was supervised and reviewed by N.N.H. All authors reviewed the manuscript.

