## Supplemental Figures for "Maternally administered naltrexone and its major active metabolite 6β-naltrexol transport across the placental barrier *in vitro*"

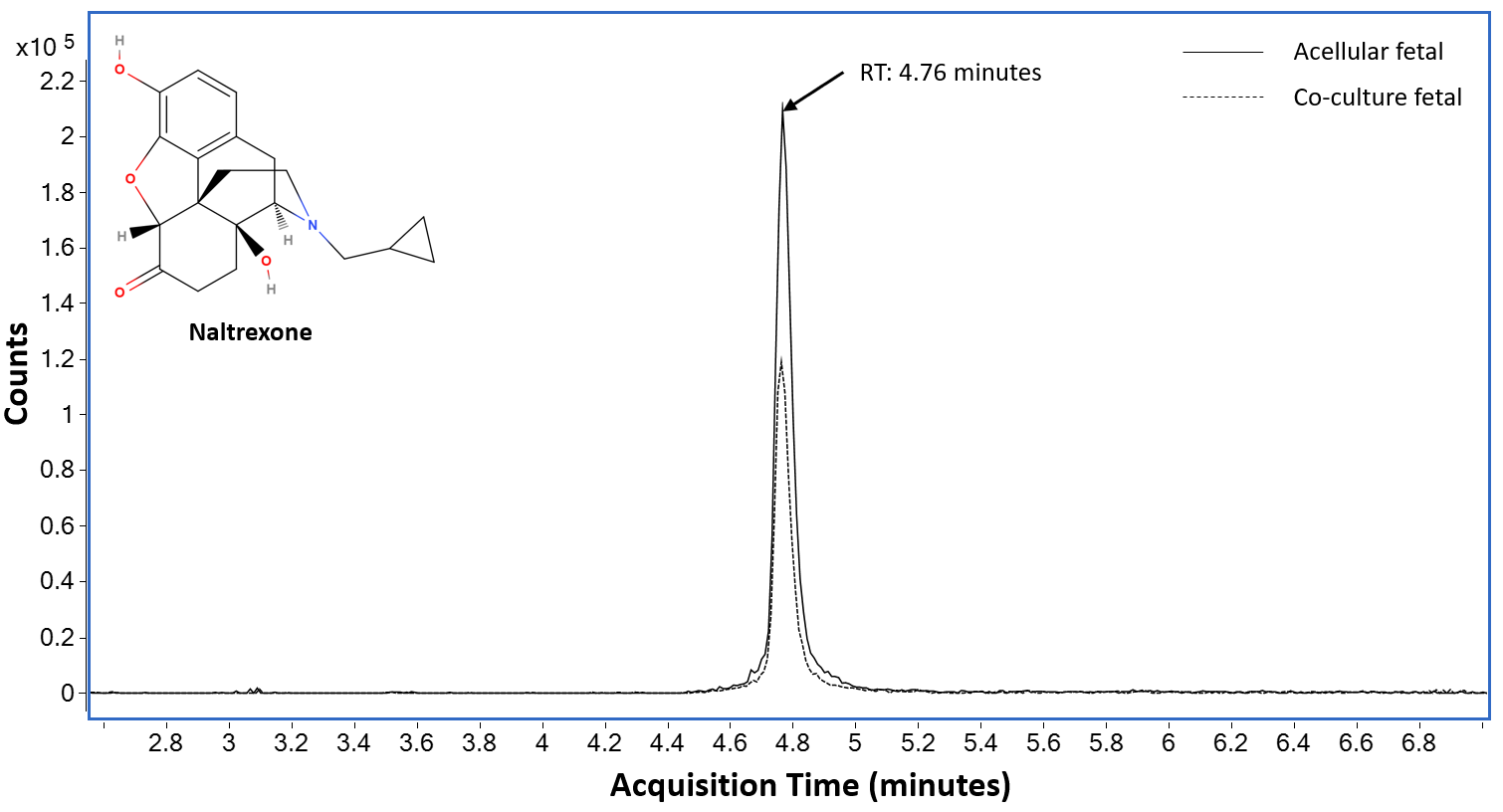

**Figure S1.** Sample Chromatogram-overlay derived from liquid chromatography/mass spectrometry (LC-MS) for the detection of naltrexone (NTX) before adjusting to the actual volume of the sample collected from the outflow of the fetal channel. Co-culture devices have both epithelial and endothelial cells and acellular devices contain only the bare membrane with perfusion of media.

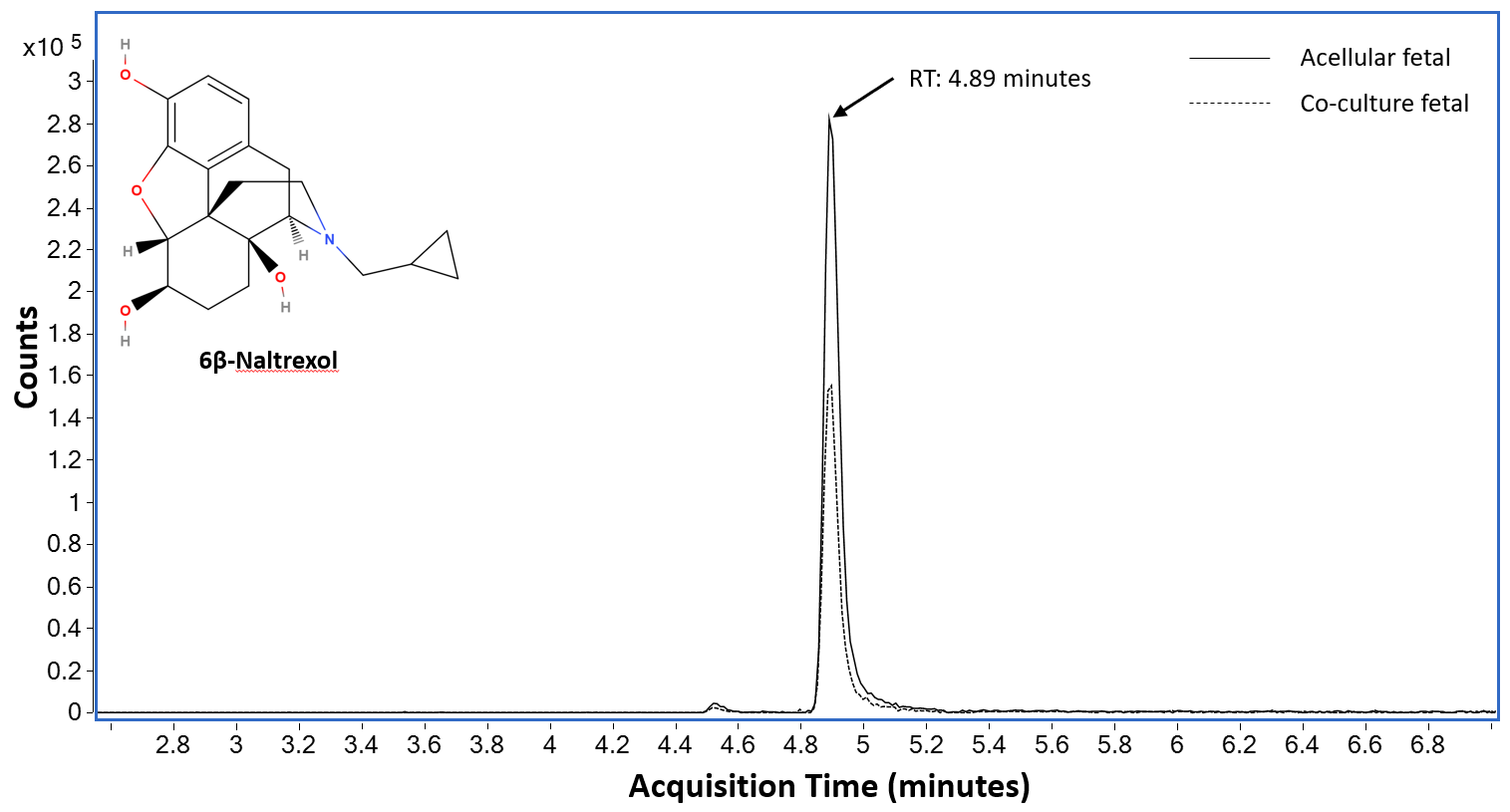

**Figure S2.** Sample Chromatogram-overlay derived from LC-MS for the detection of 6β-naltrexol before adjusting to the actual volume of the sample collected from the outflow of the fetal channel.
